# The identification of critical lethal action in antimicrobial mechanism of glycerol monomyristate against foodborne pathogens

**DOI:** 10.1101/336354

**Authors:** Song Zhang, Jian Xiong, Wenyong Lou, Zhengxiang Ning, Denghui Zhang, Jiguo Yang

## Abstract

Glycerol monomyristate (GMM) is a promising antimicrobial substance due to its broad antibacterial spectrum: however, the critical lethal action in its antimicrobial mechanism for foodborne pathogens remains unclear. In the present study, the inhibitory activities of GMM on *Escherichia coli* (*E. coli*), *Staphylococcus aureus* (*S. aureus*) and *Candida albicans* (*C. albicans*) were compared, and its membrane and intracellular action mechanism was investigated. The results showed that the susceptibility of *E. coli* to GMM was the highest, followed by *S. aureus*, and *C. albicans* being the poorest. Using flow cytometry, the GMM dose causing above 50% permeability ratio on *E. coli* was lower than that on *S. aureus*. The images from scanning electron microscope revealed no doses difference existed between the two strains when the obvious cell damage occurred. Furthermore, cell cycle and multiple fluorescent staining assays showed only the cell division of *E. coli* and *S. aureus*, excluding that of *C. albicans*, was obviously affected at 1/4 MIC and 1/2 MIC, indicating that the DNA interfere and subsequent cell division inhibition was likely to be the critical lethal action with doses near MIC, which can also explain the poor sensitivity of *C. albicans*.

**Importance:** **Foodborne** pathogens, as a common source of biological pollution in the food industry, can cause millions of food poisoning incidents each year, which poses great risks to consumers’ health and safety. The use of monoglyceride as an edible surfactant to inhibit the growth of food-borne microorganisms has been a long time, but the relevant antibacterial mechanism is too broad to accurately grasp its key lethal effect and its action doses, which not only affects the antibacterial efficiency, but also may result in the abnormalities of food flavor when adding at overdoses. The significance of the study is to identify the key lethal effect and its action doses, which will greatly enhance the understanding of the response mechanism of different types of foodborne pathogens to monoglycerides, and provide a more reasonable reference for differential control and treatment of different gastrointestinal infections when combined with antibiotics in clinical.

## Introduction

**Food** safety is very important to global public health. The control of foodborne pathogens has always been the major task in the food industry. There are millions of gastrointestinal infection cases every year due to consuming foods contaminated with harmful microbes (1). Owing to the concern of the potential toxicity of chemical preservatives (2), researchers have been striving to find effective and safe substances to replace traditional chemical preservatives such as benzoate, sorbate, and propionate (3). Monoglycerides are important non-ionic surfactants with broad antibacterial spectrum, strong antibacterial activity, and high stability (4). Among them, glycerol monomyristate (GMM) show both excellent bacterial and fungal inhibitory ability (5, 6). For example, myristic acid and its monoglyceride showed a moderate inhibition effect on the growth of *Aspergillus, Penicillium,* and *Fusarium spp.* in a reversible manner (7). An additional positive attribute of GMM is that it is digested in the gut and therefore is regarded as a safe food preservative (8).

**Monoglyceride** is often regarded as efficient antimicrobial agent, and has a high degree of membrane affinity. (9). The widely recognized action mode is that hydroxyl group in monoglyceride is adsorbed to the polar part of the cell membrane surface with the acyl carbon chain inserting into the hydrophobic region of the membrane, then moving across the phospholipids bilayers driven by the hydrophobic interaction, resulting in cell membrane perforation and the final cell death (10). It is common observed that the cell membrane is destructed and intracellular substances are released by scanning electron microscope (SEM) and other methods ^11^, ^12^. However, the critical lethal effect in the test of monoglyceride bacteriostasis is still unknown. Besides, the antibacterial performance of surfactant is based on the real exposed concentration, which is usually affected by some specific conditions, such as the barrier of biofilm, the adsorption of non-biological organics and the presence of high ion concentration (11-14). The concentration that can effectively inhibit the growth of pathogenic microorganisms is called the minimum inhibitory concentration (MIC), which also provides us an effective way to identify the critical lethal effect.

**According** to some previous reports, foodborne pathogens can be divided into gram-negative bacteria (*Escherichia coli, Salmonella typhimurium, Vibrio parahaemoliticus, Clostridium perfringens, Klebsiella pneumoniae*), gram-positive bacteria (*Staphylococcus aureus, Listeria monocytogenes, Clostridium botulinum, Bacillus cereus, Bacillus anthracis*), and fungi (*Candida albicans, Candida parapsilosis, Epidermophyton floccosum, Trichophyton mentagrophytes, Trichophyton rubrum)* (1, 8, 15, 16). Therefore, *E. coli, S. aureus,* and *C. albicans* were selected as model microorganisms representing gram-negative and gram-positive bacteria, as well as fungi to complete this research.

**This** study was designed to compare the sensitivity of *E. coli, S. aureus,* and *C. albicans* to GMM and investigated the action mode at different exposed doses in order to search the critical lethal effect in antimicrobial test, which had not been reported previously. The antibacterial curves were employed to study the cell viability after adding GMM. The changes in membrane morphology and permeability ratio were assessed SEM and flow cytometry. In addition, the impact of GMM on DNA double helix and cell division was assessed by UV-visible absorption spectrum and cell cycle assay. Finally, the synthesis inhibition of DNA, RNA, and protein was measured to explain the potential relationship between DNA double helix damage and cell division inhibition. With further understanding of antibacterial mechanisms, it is demonstrated that DNA interference and subsequent inhibition of cell division are the key lethal effect in GMM assay, which also provides a guide for the design of antibiotics and the control of drug-resistance bacteria.

## Results

### Sensitivity of *E. coli, S. aureus, and C. albicans* to GMM

**The** *E. coli, S. aureus and C. albicans* cells were effectively inhibited with GMM at 32, 64 and 500 μg/mL, respectively, which indicated their corresponding MIC values. As shown in Figure 1A, the growth of viable *E. coli, S. aureus, and C. albicans* was inhibited by GMM at various degrees. Specifically, the growth of *E. coli* and *S. aureus* was totally inhibited at 125 μg/mL and 500 μg/mL, respectively, whereas *C. albicans* cells could not be completely inhibited until a concentration of 1,000 μg/mL was used, suggesting that the sensitivity of *E. coli* to GMM were the strongest, followed by *S. aureus*, and that of *C. albicans* was the weakest.

**Figure 1.**
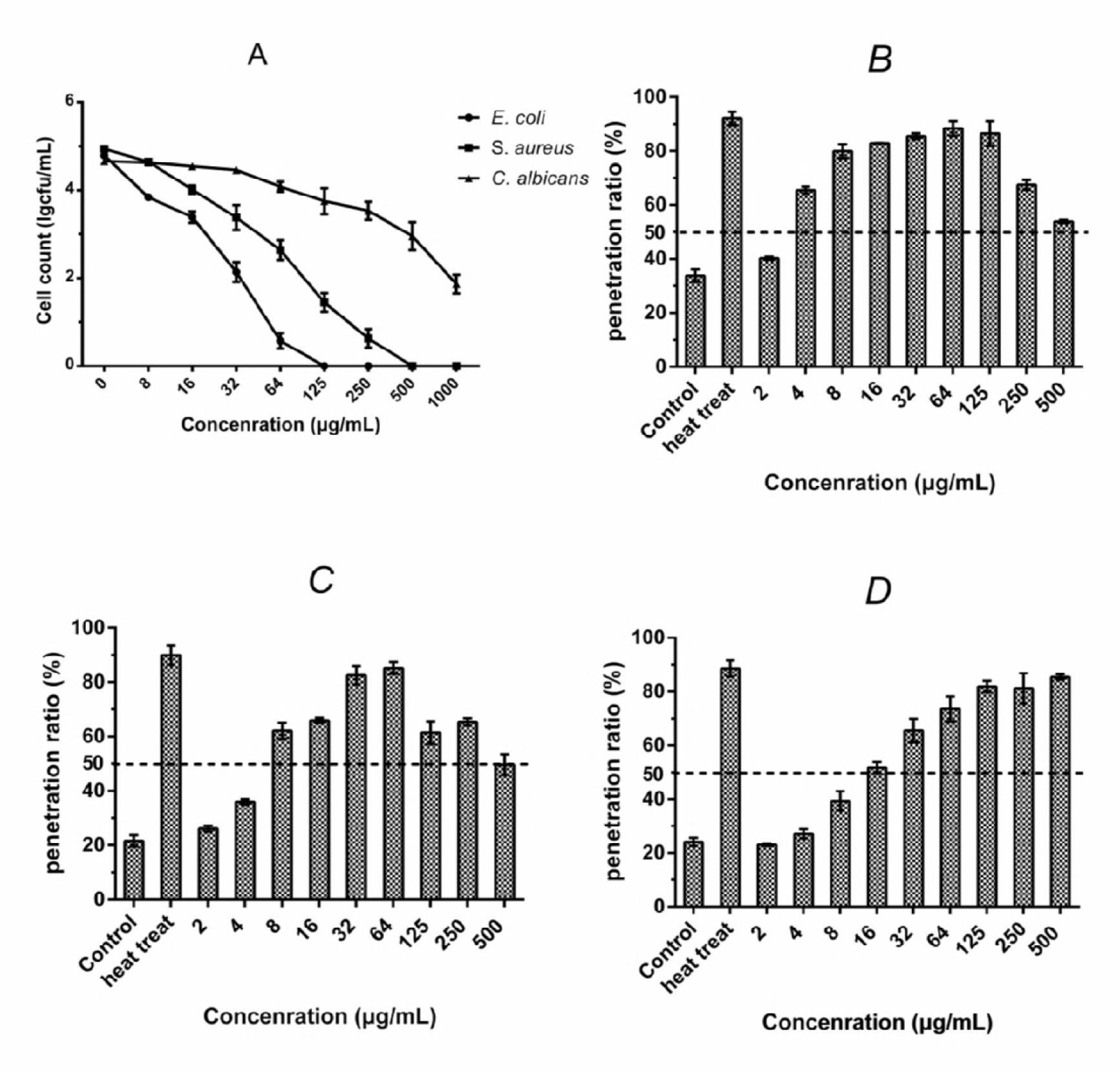
Cell count of viable *E. coli, S. aureus,* and *C. albicans* after treatment with GMM at 0, 8, 16, 32, 64, 125, 250, 500, and 1000 μg/mL for 1 h was shown (A). Penetration ratio of *E. coli* (B), *S. aureus* (C), *and C. albicans* (D) treated with GMM at concentrations ranged from 2 to 500 μg/mL. Error bars in histograms represent standard deviations in triplicate. The dotted line in the figure represented the 50% level in penetration ratio.

### Permeability effect of GMM on microbial cells

**As** shown in Figure 1B-D, the proportion of three pathogens cells with damaged membranes all increased with raising GMM concentration to MIC, but did not exceed the corresponding MDB. Specifically, the permeability ratio in *S. aureus* and *C. albicans* groups both showed evidently growth trend instead of that in *E. coli* group, which remained above 60% of penetration ratio in the range of 4 μg/mL to 500 μg/mL. In terms of the membrane penetration level, the concentrations caused above 50% penetration ratio in *E. coli, S. aureus, and C. albicans* assays were 4 μg/mL, 8 μg/mL, and 16 μg/mL, respectively. Surprisingly, increasing further GMM concentration to 500 μg/mL resulted in a decrease in permeability ratio of *E. coli* and *S. aureus*, which could be explained by detection deviation caused by a large proportion of membrane lysis and cell death under high exposure concentration (17).

### Cell morphology changes induced by GMM

**As** shown in Figure 2 A0 to C0, the *E. coli, S. aureus, and C. albicans* in the control groups all had completely flat surfaces without any defects. No influence was seen on *E. coli* cell membranes after treatment with GMM at 1/2 MIC, whereas rough and recessed surfaces appeared successively on *E. coli* cells exposed to GMM at 1 MIC and 2 MIC (Figure 2 A1–A3). Unlike *E. coli, S. aureus* surfaces appeared slightly rough at 1/2 MIC (Figure 2 B1). After treatment with GMM at MIC, the original rough surface developed into a wrinkled and concave structure, and increasing the concentration further to 2 MIC led to breakage and lysis appearing on cell membranes, which suggested the destruction level of GMM on *S. aureus* cell membrane was greater than that of GMM on *E. coli at* 2 MIC (Figure 2 B2 and B3). Examination of *C. albicans* showed increasing rough and wrinkled changes on cell membranes when GMM concentration was increased from 1/2 MIC to 2 MIC, but breakage and defection was not observed until the final concentration (Figure 3 C1-C3).

**Figure 2.**
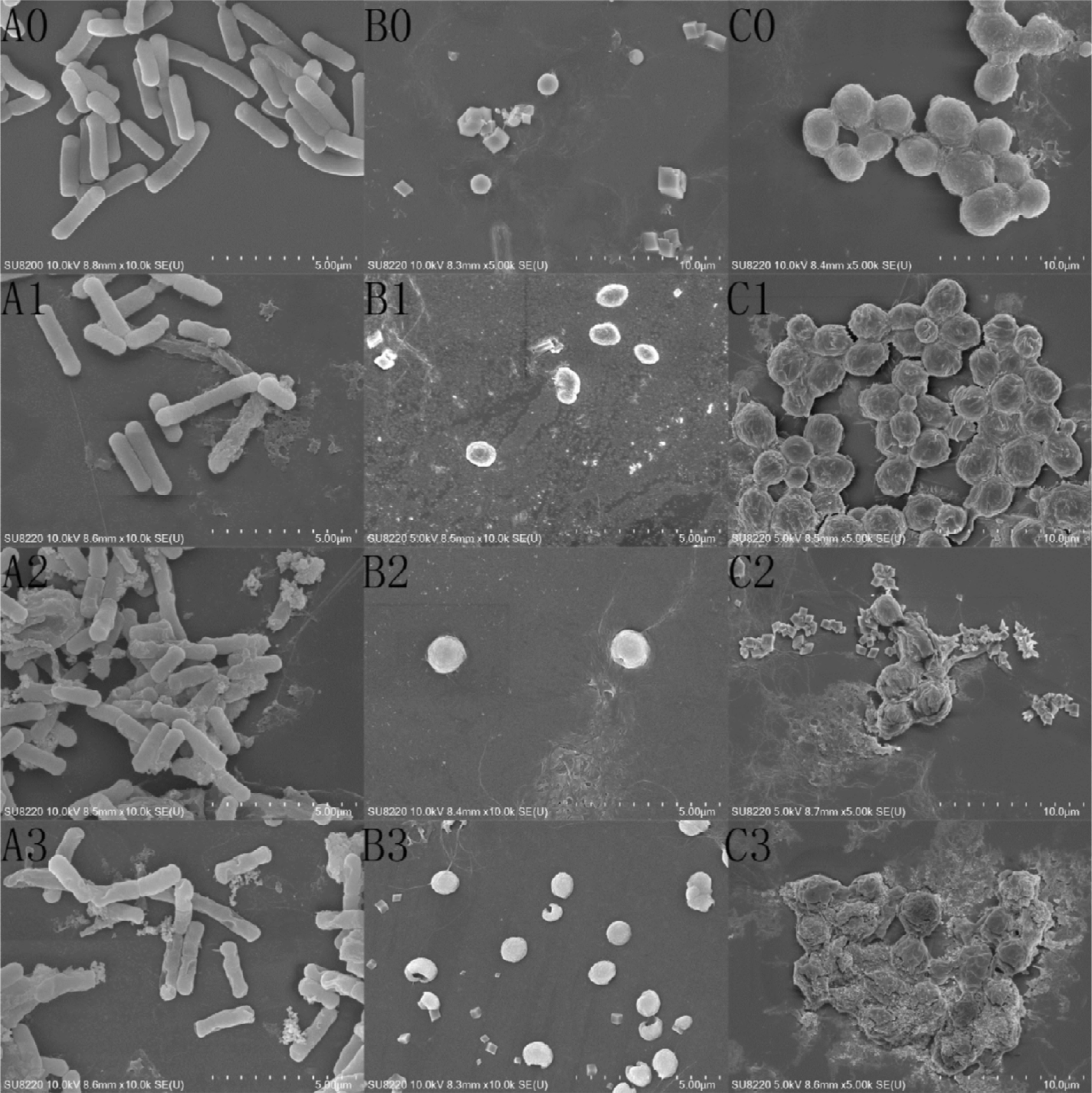
SEM images of *E. coli* (A0-A3), *S. aureus* (B0-B3), *and C. albicans* (C0-C3) treated with GMM at 0 (A0, B0, and C0), 1/2 MIC (A1, B1, and C1), 1 MIC (A2, B2, and C2), and 2 MIC (A3, B3, and C3) was recorded. No monoglycerides were added to the control groups (A0, B0, and C0). Three images of each cell sample were recorded under the magnification of 10,000 times for *E. coli* and *S. aureus*, and 5000 times for *C. albicans.*

**Figure 3.**
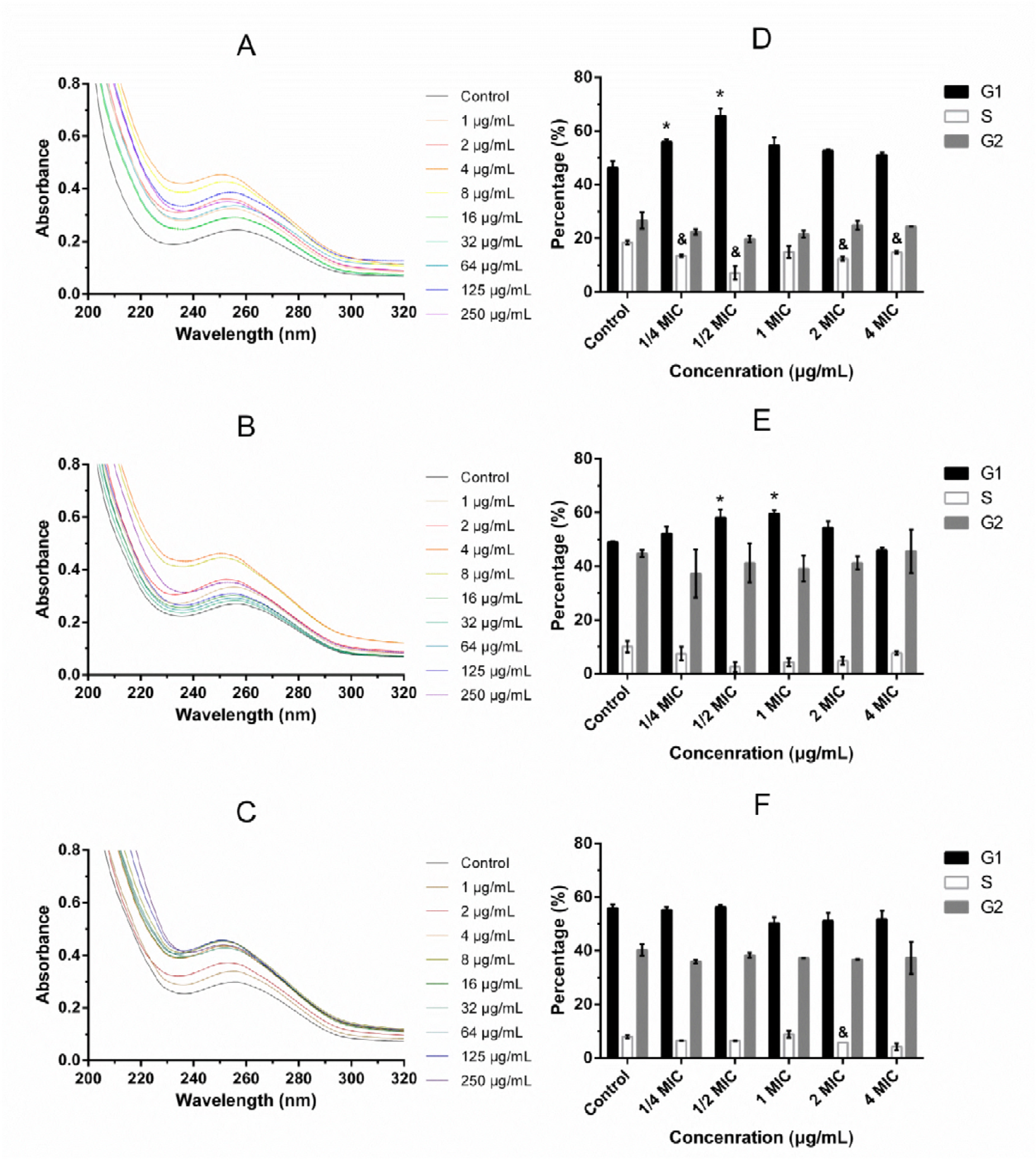
The UV spectrums of genomic DNAs from *E. coli* (A), *S. aureus* (B), *and C. albicans* (C) cells treated at 1, 2, 4, 8, 16, 32, 64, 125, and 250 μg/mL, and percentages of G1, S, and G2 phases in *E. coli* (D), *S. aureus* (E), *and C. albicans* (F) after adding GMM with concentrations at 1/4 MIC, 1/2 MIC, 1 MIC, 2 MIC, and 4 MIC were measured. All data were the average of three determinations in parallel and error bars represented standard deviations. “*, & and #” indicated statistical significant variance from the control groups in G1, S, and G2 phases, respectively, if p<0.05.

### Interaction between GMM and genomic DNA of pathogenic microbial cells

**As** shown in Figure 3A-C, the absorbance peak value of genomic DNA from *E .coli, S. aureus, and C. albicans* cells at 260 nm firstly increased with a slight blue shift in wavelength (from 257 nm to 250 nm for *E. coli,* from 256 nm to 251 nm for *S. aureus* and from 257 nm to 250 nm for *C. albicans*) with increasing GMM concentrations, which is called the DNA hyperchromic effect. However, further increasing GMM concentration caused a clear decrease in OD_260_ accompanied with a slight red shift in wavelength (from 250 nm to 256 nm for *E. coli,* from 251 nm to 257 nm for *S. aureus* and from 250 nm to 255 nm for *C. albicans*). Examination of the differences in Figure 3A–C, we found that the GMM concentration causing the maximum absorption peak of *C. albicans* DNA at 260 nm was significantly greater than that inducing the maximum absorption peak of *E. coli* and *S. aureus* DNA at 260 nm (with corresponding GMM concentrations of 16, 4, and 4 μg/mL).

**Interference of GMM on cell cycle**

**The** *E. coli* and *S. aureus* both belong to bacterial cell that has a cell cycle with I, R, and D phases, which corresponded to the G0/G1, S, and G2/D phases in eukaryotic cell such as *C. albicans*. As shown in Figure 4, the peak shape of the flow histograms of *E. coli* cells first widened and then narrowed in dose range from 1/4 MIC to 4 MIC, whereas the flow peak shape of *S. aureus* and *C. albicans* changed little. The specific changes of the G1, S, and G2 phase proportion of microbial cells after GMM treatment are shown in Figure 3D-F. The G1 ratio in *E. coli* and *S. aureus* groups increased to varying degrees with increasing GMM concentration, and the significant effect was observed at 1/4 MIC and 1/2 MIC for *E. coli*, 1/2 MIC and 1 MIC for *S. aureus*. Surprisingly, no significant increase or decrease appeared in the G1 ratio of *C. albicans* with GMM doses increased from 1/4 MIC to 4 MIC, suggesting that the cell cycle of *C. albicans* was less likely to be disturbed in GMM treatment. In addition, further increasing GMM doses to 4 MIC led to a decrease of G1 ratio in *E. coli* and *S. aureus*, causing by membrane damage and DNA leakage under high exposure doses, which could be demonstrated by the measurement in Figure 1 and the observation in Figure 2. Examination the differentiated performances of the cell cycle among three pathogens showed that the interference level of GMM on *E. coli* cell cycle was greater than that on *S. aureus*, and no any disturbance was observed against *C. albicans* cell cycle.

**Figure 4.**
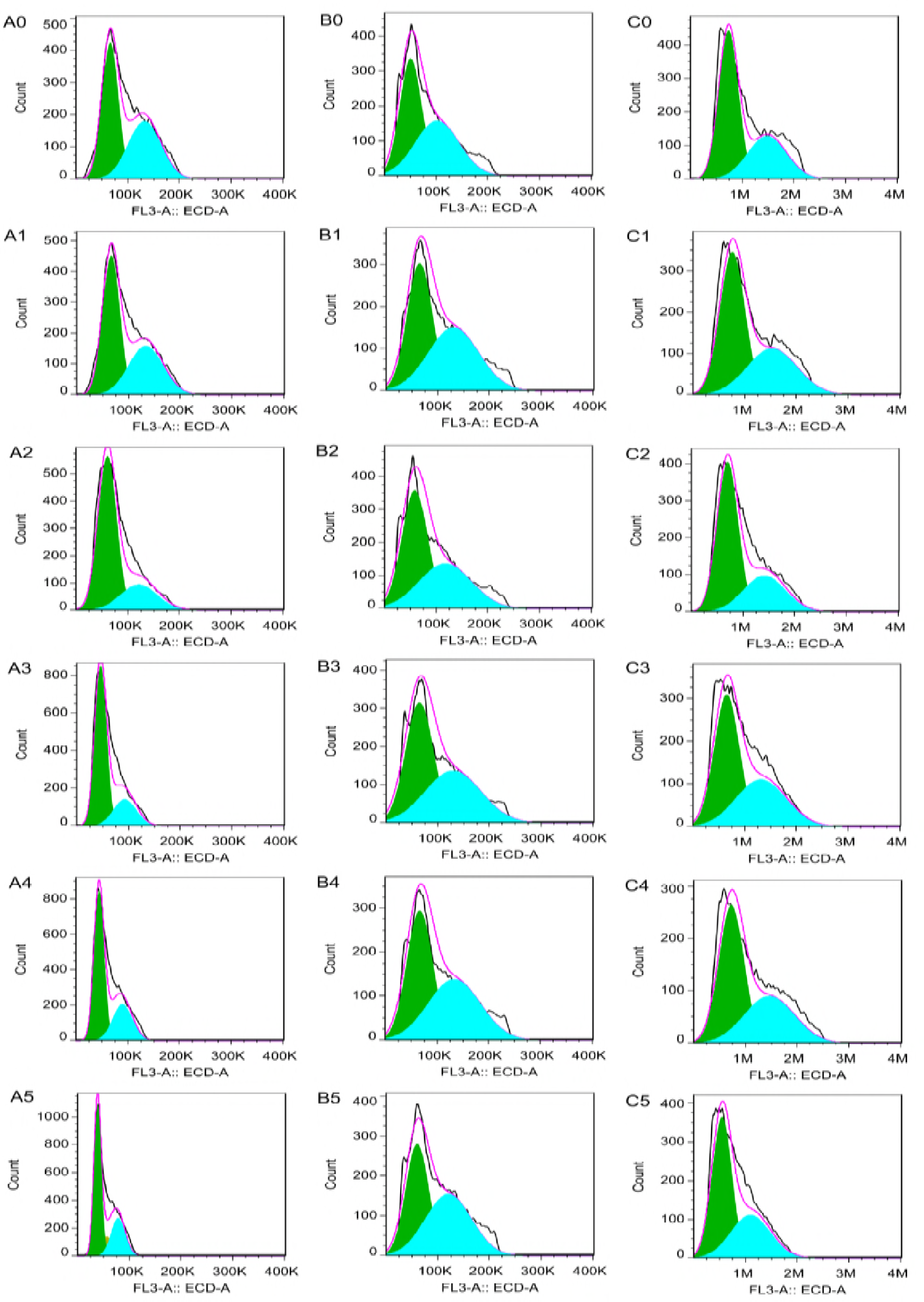
Flow histograms of *E. coli* (A0-A5), *S. aureus* (B0-B5), *and C. albicans* (C0-C5) treated with GMM at 0 (A0, B0, and C0), 1/4 MIC (A1, B1, and C1), 1/2 MIC (A2, B2, and C2), 1 MIC (A3, B3, and C3), 2 MIC (A4, B4, and C4), and 4 MIC (A5, B5,and C5) was shown.

### Inhibition of GMM on intracellular DNA, RNA, and protein synthesis

**In** addition to the study on the effect of GMM on the structure and function of bacteria genomic DNA, the changes of DNA, RNA, and protein content in cell was shown in Figure 5. Compared to the steady growth of DNA, RNA, and protein fluorescence intensity in three control groups, the three biomacromolecules in treated group showed a completely different performance. Specifically, the RNA fluorescence density in *E. coli* and *S. aureus* decreased immediately without any delay on time once adding GMM, unlike to a strange trend of increasing firstly and then declining in DNA and protein light intensity, indicating that RNA synthesis was firstly disturbed, which was 30min or one generation earlier than the time when DNA and protein synthesis received inhibition. The inconsistency existed on the time point when three macromolecules received affection also suggested RNA synthesis, instead of protein and DNA synthesis, was the primary action goal in the process of cell cycle arrest. As for *C. albicans*, the light density of three macromolecules did not appear obviously change except for a decrease of RNA fluorescence density occurring at 40 min after adding monoglyceride, which was attributed to the high permeability at 1/2 MIC. Comparison the differentiated performance in DNA, RNA, and protein content among three pathogens revealed that the timely interference for RNA synthesis and delayed suppression for DNA and protein synthesis might contribute to the disturbance in cell cycle.

**Figure 5.**
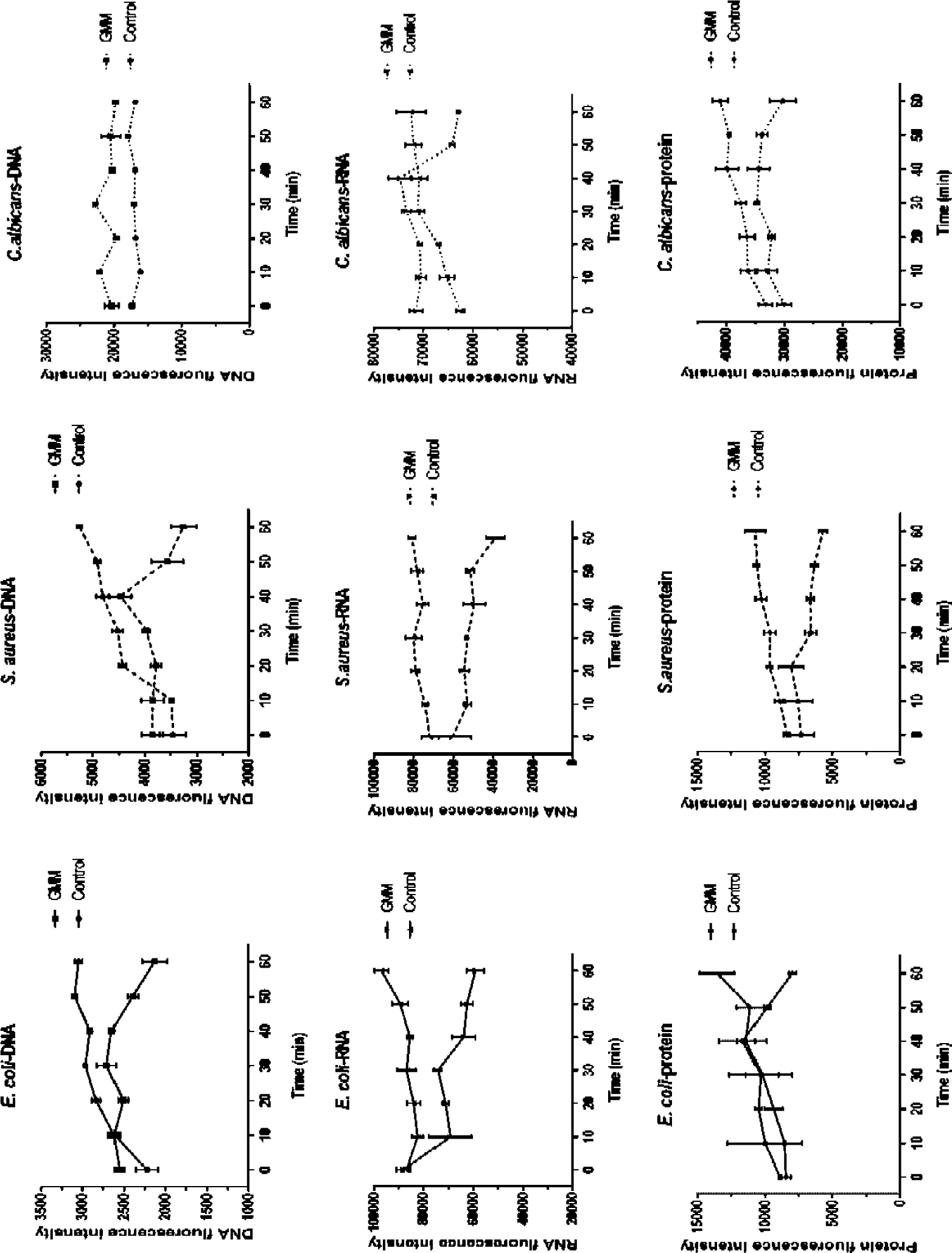
Changes of DNA, RNA, and protein fluorescence intensity in *E. coli, S. aureus and C. albicans* cells treated with GMM at 1/2 MIC was detected. The three control groups added the same volume of ethanol instead of monoglyceride solution. All data were the average of three measurements in parallel, and error bars represented the standard deviations.

## Discussion

### Sensitivity of *E. coli, S. aureus, and C. albicans* to GMM

**The** results in this assay demonstrated that GMM effectively inhibited the growth and cell viability of *E. coli, S. aureus, and C. albicans.* The MIC values of *E. coli* and *S. aureus* were similar to what has been reported previously (1, 15). The only difference we observed was that the MIC of *E. coli* was lower than that of *S. aureus*, which was a real uncommon phenomenon. A possible explanation was proposed that the lipopolysaccharide (LPS) on the surface of gram-negative bacteria increase the affinity between monoglyceride and bacterial cell, making it easier for acyl chain to enter the phospholipids bilayers (18-20). The huge difference among the MIC values of *E. coli, S. aureus,* and *C. albicans* also illustrated that the susceptibility of gram-negative, gram-positive bacteria, and yeast cells to GMM was quite different. The discrepancy may be attributed to cell membrane composition and structure, such as differences in lipid composition (21). There was evidence that the yeast cell membrane as rearranged, and the adsorbed monocaprylate was removed during the buffer wash using *E. coli* polar lipid extract and *Saccharomyces cerevisiae* polar lipid extract to form supported lipid bilayers of bacteria and yeast, respectively (18). In addition, the cell number of viable *E. coli, S. aureus and C. albicans* decreased more than 2 log units when adding GMM at 1 MIC, which was stronger than the action effect of LC_50,_ suggesting that the growth inhibition in three pathogens was closely related to the loss of cell viability in GMM treatment.

### GMM penetrates and destroys cell membranes

**Many** studies indicated that cell membrane was the primary target of action, as monoglyceride was amphipathic substance which could interact with and destabilize cell membrane (19, 20, 22). SEM images indicated smooth and intact cell membrane of *E. coli, S. aureus and C. albicans* was disturbed and destroyed after adding GMM, which was related to the incorporation of GMM into the membrane (23) and the decrease of ability to respond to external stress (24). The action dose was usually 2 MIC or more than 2 MIC when membrane breakage or cell lysis of three pathogens occurred. The integration of GMM into the microbial cell surface also led to changes in cell membrane permeability. Some studies reported GMM diffused through the cell outer membrane and the cell wall creating traversable holes, resulting in the loss of membrane fundamental function and an obvious increase in cell permeability (1, 19, 25). The results in the flow cytometry assay suggested a positive correlation between greater cell membrane permeability and higher GMM concentration, except for *E. coli* and *S. aureus* whose penetration ratios even showed a decline when further increasing to a much higher dose than the MIC. In addition, Bunkova also found that monoglyceride displayed a membrane damage level against pathogens in a dose-dependent manner, which was consistent with the above observation (26).

**It** is necessary to further consider the correlation between MIC, the cell morphology change, and membrane permeability increase for finding the critical lethal action and comprehensive understanding of the reaction mechanism to external surfactant. As shown in Figure 1B-D, the GMM concentration causing more than 50% maximal permeability ratio was 4, 8, and 16 μg/mL for *E. coli, S. aureus and C. albicans,* respectively, which were far lower than their respective MIC. However, the above doses did not result in a significant decrease in cell count, indicating that membrane permeability increase did not cause the obvious loss of cell viability of pathogens. Similarly, a remarkable depression and breakage appeared at 2 MIC (64 μg/mL) for *E. coli*, 1 MIC (64 μg/mL) for *S. aureus*, and more than 2 MIC (exceed 1000 μg/mL) for *C. albicans*, which was more like the action result of antibacterial agents. All the above observations suggested that GMM increase membrane permeability at low doses and induce membrane damage or even cell lysis at high doses above MIC. What’s more, the MIC of *C. albicans* was almost 32 times of its concentration causing above 50% permeability, which was far higher than the 8 times in *E. coli* and *S. aureus*, implying that there may be potential intracellular action goals to be identified after GMM crossing cell membrane.

### Interference of GMM on genomic DNA

**Using** the UV-visible absorption spectrum was an effective way to investigate the interaction between monoglycerides and genomic DNA in microbial cells (27). It was reported that DNA had a strong absorption peak at 260 nm, which was derived from the strong absorption of purine and pyrimidine bases in DNA, and the location and intensity of the DNA absorption peak could be migrated and changed when a foreign substance was bound to DNA (28, 29). In the presence of GMM, the absorption peak of DNA at 260 nm increased gradually with a slight blue shift as increasing GMM concentration, which is called hyperchromic effect (30). This type of change in DNA spectra has been regarded as a symbol of the destruction of the double helix in genomic DNA in the case of antibacterial ingredients binding to DNA (31). In contrast to previous studies, further increasing GMM concentration resulted in a gradual loss of the DNA hyperchromic effect, and the cause for this uncommon change was still unknown. The nucleic acid structure changes caused by GMM might affect normal cell function, which needs to be further investigated in the future (32).

**Cell** cycle was an important indicator of cell division function, which was reflected as cell proportions at different stages of cell division. Flow cytometry results indicated that GMM disrupted phase G1 instead of phases S and M, causing microbial cells to stay at phase G1 and the ultimate termination of the cell cycle. The disruption of GMM on the DNA double helix could inhibit the synthesis of essential materials for DNA replication (33). As shown in Figure 3, GMM interfered cell cycle in a dose-dependent manner, which could be described as a more obvious interruption effect with increasing GMM concentration at a low dose range, and further increases led to a diminished effect. The changing trend was related to the penetration efficiency of GMM on different microbial cell membranes and supersaturated DNA binding sites (34). As for *C. albicans,* no significant increase in G1 phase was observed, which might due to the protection of nuclear membrane to genomic DNA. According to the reports on the correlation between intracellular action targets and antimicrobial activity in recent reports, it was speculated that GMM might exert its antimicrobial activity by destroying the DNA double helix and affecting normal cell division (35-37).

**To** establish a relationship between DNA structure interference and cell cycle change, the intracellular biological macromolecule measurement was performed using HO, PY, and FITC for staining DNA, RNA, and protein, respectively. This multi-fluorescence measurement effectively characterized the overall imbalance conditions of DNA, RNA, and protein caused by cell cycle-interfering agent (38). In the current study, the time point when DNA and protein received interference in *E. coli* and *S. aureus* was clearly later than that of RNA, whose content declined immediately after adding GMM. The 30-minute time interval implied that the transcription of DNA to RNA was the primary inhibitory process in the interaction between GMM and genomic DNA. In addition, the smooth growth of DNA and protein content, as well as the slight drop in RNA content, suggested that the synthesis of three major macromolecules in yeast has not received significant impact except that a small amount of RNA leaked to the extracellular, which contributed to demonstrate that the cell division of *C. albicans* was not affected by GMM treatment. Combined with previous findings (39, 40), a potential DNA inhibitory route was proposed in which DNA transcription was first suppressed due to the destruction of the double helix caused by GMM. Then, the translation process of mRNA into proteins, including synthesis of enzymes related to DNA replication, was affected, thus DNA replication was postponed and the cell cycle was blocked in phase G1, and finally resulting in cell division disorder.

**A** new concentration-dependent antibacterial mechanism of GMM on *E. coli* and *S. aureus* were summarized and presented in Figure 6. The glycerol mono-fatty acid ester with 14-carbon acyl chain has a membrane permeability of 50% or more at 1/8 MIC. After penetrating through cell membrane, GMM acts on DNA transcription, leading to DNA replication restriction and subsequent cell cycle arrest, ultimately inhibiting bacterial cell division. This type of intracellular action usually occurs at concentrations close to the MIC, thus it is regarded as the critical lethal effect in the test of GMM antibiosis. And at higher exposure doses, like 2 MIC or more, cell membrane damage or even cell lysis will occur, which is more like the result of a combined action of many antibacterial effects.

**Figure 6.**
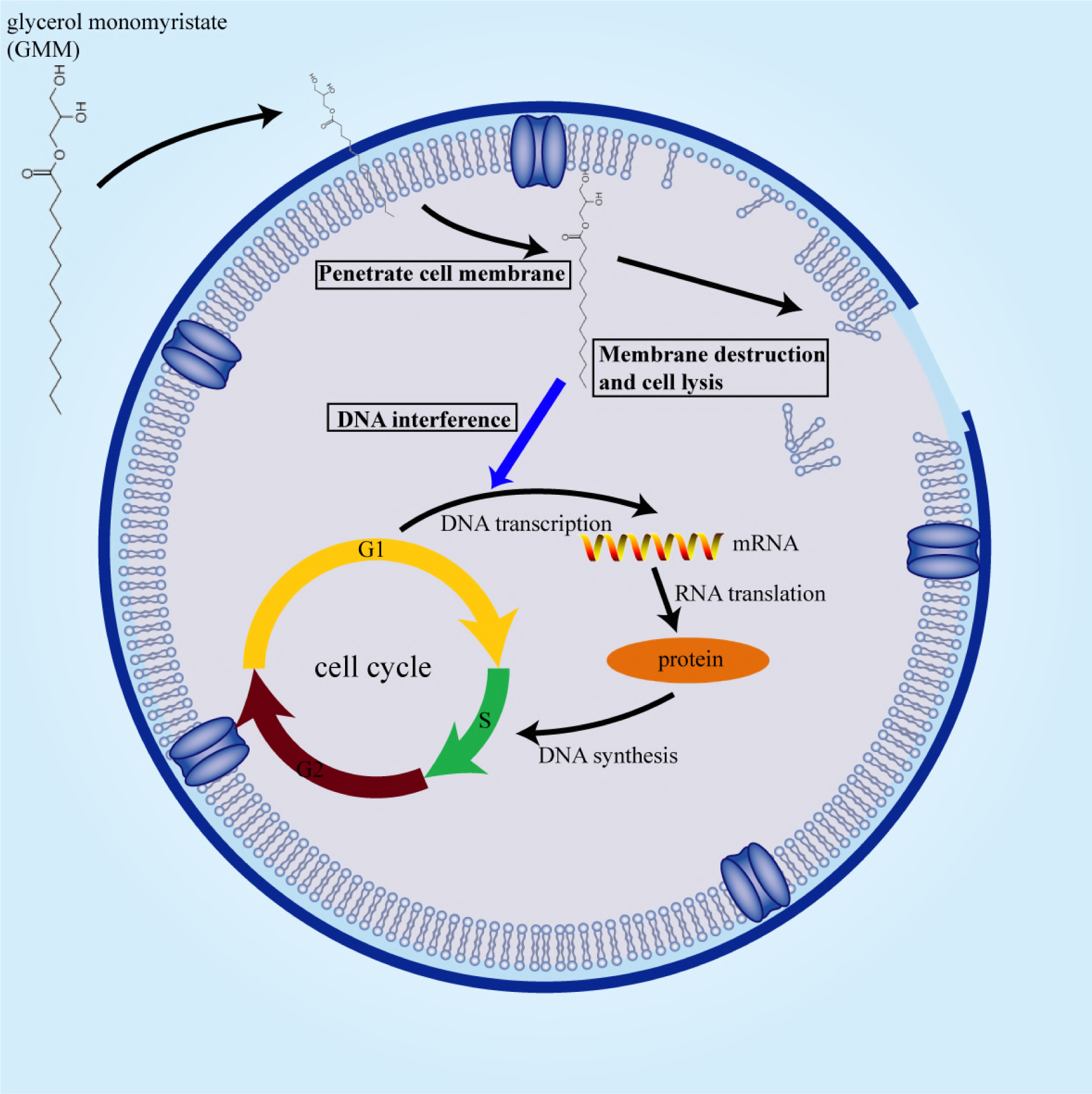
Membrane and intracellular action mechanism of GMM was shown. GMM firstly crossed the cell membrane and interfered with the normal function of the DNA, eventually leading to cell lysis. The action site of GMM on DNA was identified as the process of DNA transcription, causing the reduction in the synthesis of RNA and protein, resulting in cell cycle arrest and ultimately cell division inhibition.

**However**, *C. albicans* did not appear the similar experimental expectations as bacteria in cell cycle and biomacromolecules measurement assays. Although the double helix of yeast genomic DNA extracted in vitro may be destroyed by GMM, biological experiments in medium have shown that the cell cycle of *C. albicans* has not received significant interference, and intracellular biomacromolecules synthesis has also not been obviously affection, indicating that GMM has no inhibitory effect on cell division of *C. albicans*. Taking into account the fact that the MIC of *C. albicans* is 32 times the concentration of membrane permeability change, totally different from that in bacteria, thus we have reason to believe that the DNA interfere plays a key role in the sensitivity divergence. On the whole, the antibacterial mechanism can be classified as follows: increase of membrane permeability is the basis of antibacterial action, cell division inhibition is the critical lethal effect, and cell lysis is the result of many antibacterial actions.

**In** summary, the present study compared the sensitivity of *E. coli, S. aureus,* and *C. albicans* to GMM, and explained the causes for the variance in sensitivity from the aspects of membrane permeability increasing, cell division inhibition and cell lysis. The antimicrobial assay indicated that *E. coli* was the most sensitive to GMM treatment, followed by *S. aureus*, and the worst sensitivity was belong to *C. albicans*. The higher membrane permeability at low concentrations demonstrated that the sensitivity of *E. coli* was indeed better than that of *S. aureus.* As for SEM experiment, the changes of the cell membrane morphology between the two pathogens were similar at the same concentration. Furthermore, UV-visible spectrum, cell cycle and three biomacromolecules detection were conducted to make clear the reason for the sensitivity discrepancy between bacterial and yeast to GMM. Although the double helixes of the extracted DNA from three pathogens were damaged, only *E. coli* and *S. aureus*, excluding *C. albicans*, showed cell division inhibition in GMM treatment. The correlation between loss of DNA suppression mechanism and poor sensitivity in *C. albicans* confirmed cell division disorder was the critical lethal effect.

## Materials and methods

### Chemicals and agents

**GMM** (purity≥99%) was purchased from Molbase Chemical Co. (Shanghai, China). A series of GMM stock solutions were prepared by dissolving it in absolute ethanol to obtain concentrations of 0.01, 0.02, 0.04, 0.08, 0.16, 0.32, 0.64, 1.25, 2.5, 5, and 10 mg/mL. Propidium iodide (PI, purity≥95%) was obtained from Aladdin Biochemical Technology Co. (Shanghai, China). A 10 mg/mL PI stock solution was made in phosphate buffer solution (PBS, pH 7.2) and stored at −4°C for later dilution use. An Ezup column bacteria genomic DNA purification kit and an ezup column yeast genomic DNA purification kit were both bought from Sangon Biotech Co. (Shanghai, China). Tris-HCl buffer (0.05 M, pH 7.4) was obtained from Yuan Ye Biological Technology Co. (Shanghai, China). Hoechst 33342 (HO, purity≥98%), Pyronin Y (PY, purity≥75%), and fluorescein isothiocyanate (FITC, purity≥95%) were purchased from Yuan Ye Biological Technology Co. (Shanghai, China). The stock solutions of three dyes, with concentrations of 500 μg/mL of HO, 2 mg/mL of PY and 100 μg/mL of FITC, were prepared in PBS and refrigerated for later use.

### Strain activation and culture

**Three** foodborne pathogenic microorganisms, *Escherichia coli O157:H7 ATCC35150, Staphylococcus aureus ATCC25923* and *Candida albicans ATCC10231* were purchased from Guangdong Culture Collection Center (Guangzhou, China). The strains were activated by dissolving them in 1 mL sterile PBS prior to transferring onto a solid plate medium containing either tryptone soy agar (TSA, bacterial solid medium) or Sabouraud dextrose agar (SDA, fungal solid medium) and incubated for 24 h at 37°C (48 h at 28°C for *C. albicans*). A loop of a single colony from the above solid plate media was then inoculated onto a TSA or SDA slope with multiple cross operations and cultured for 24 h at 37°C (48 h at 28°C for *C. albicans*) followed by refrigerated store at 4°C.

### MIC determination

**The** refrigerated strains were cultivated in 100 mL tryptic soy broth (TSB, bacterial liquid medium) or Sabouraud dextrose broth (SDB, fungal liquid medium) with orbital shaking at 120 rpm in 37°C (28°C for *C. albicans*) until the mid-logarithmic period was achieved. The cell collection was conducted by centrifugation at 3000 rpm for 5 min followed by washing twice with sterile PBS. The cell pellets were then resuspended with sterile broth to the optical density of 0.05 at 600 nm (OD_600_≈ 0.05). The subsequent experimental operation referred to Byeon’s method in a 96-well microtiter plate (41). The MIC was defined as the lowest GMM concentration which prevented bacteria growth for 24 h (48 h for *C. albicans*) when the absolute value of the difference between the initial and final OD_600_ was less than 0.05.

### Antibacterial curve assay

**The** microbial cell populations after GMM treatment were measured by the plate count method according to a previous study (42). The pathogen cells were collected as described above and resuspended to a final cell concentration of approximate 10^5^ CFU/mL (OD_600_≈ 0.05) with sterile broth. Culture solutions (950 μL) of *E. coli, S. aureus*, and *C. albicans* were mixed with 50 μL different concentrations of GMM solution to achieve final monoglyceride concentrations of 8, 16, 32, 64, 125, 250, 500, and 1000 μg/mL. For the controls, 50ul ethanol was added instead of GMM. The experiment was performed in triplicate. All sample tubes were cultivated for 1 h at 37°C (2 h at 28°C for *C. albicans*) with orbital shaking at 120 rpm. Subsequently, a 10-fold serially dilution was performed on the samples followed by culturing and subsequent counting of the colony forming units (CFUs).

### Cell permeability test

**The** changes in the membrane permeabilities of *E. coli, S. aureus, and C. albicans* cells treated with GMM were assessed by using flow cytometry (43). The 950 μL cell suspensions (cell density≈ 10^5^ CFU/mL) were treated with 50 μL different concentrations of GMM to achieve final concentrations of 0, 1/4 MIC, 1/2 MIC, 1 MIC, 2 MIC, and 4 MIC before incubation for 1 h at 37°C (2 h at 28°C for *C. albicans*). For the negative control groups, 50 μL of ethanol was added instead of GMM, whereas the cells in the positive control groups were treated with constant temperature heating for 1 h at 85°C in order to determine the maximum detection boundary. All samples including the experimental and control groups were performed in triplicate. Subsequently, the microbial cells were collected by centrifugation and resuspended in 1 mL sterile PBS followed by staining with 50 μL of 100 μg/mL PI work solution. The stained cells were incubated for 20 min at room temperature in dark conditions followed by flow cytometry analysis.

**A** semi-automatic flow cytometer (CytoFLEX, Beckman Coulter Co., CA, USA) was used to perform cell membrane permeability detection. PI dye was excited by an argon ion laser at 488 nm, and the red fluorescence signals emitted from PI-DNA were captured in the ECD channel (610/20). The sample flow rate was adjusted to 500 cells/s, and ultimately, a minimum of 10,000 cells were collected for data analysis.

### Scanning electron microscope

**A** SEM assay was performed using Marounek, et al.’s method with slight changes (9). Firstly, 950 μL aliquot cell solutions (cell density≈ 10^5^ CFU/mL) were prepared and mixed with 50 μL of different concentrations of GMM to achieve final concentrations of 0, 1/2 MIC, 1 MIC and 2 MIC. The same amount of ethanol was added to the control groups. All cell solutions were cultured for 1 h at 37°C (2 h at 28°C for *C. albicans*) prior to centrifugation and gentle washing with PBS twice. Secondly, the cells were fixed with 0.5 mL of 2.5% (v/v) glutaraldehyde in PBS overnight at 4°C and post fixed with 0.1 mL 2% (w/v) osmium tetroxide in PBS for 2 h at room temperature. Subsequently, the immobilized cells were washed with ultrapure water (18.2 MΩ cm) and dehydrated for 10 min with 30%, 50%, 70%, 90%, and 100% ethanol, respectively. The dehydrated cell samples were then resuspended in absolute ethanol and settled by dropper onto a coverslip to stand for 15min. Finally, all samples were subjected to freeze-drying under vacuum and sputter-coated prior to microscopic observation with a cold field scanning electron microscope (UHR FE-SEM SU8220, Hitachi Ltd., Tokyo, Japan).

### Interaction of GMM and genomic DNA of microbial cells

**The** genomic DNA derived from viable *E. coli, S. aureus,* and *C. albicans* cells in the mid-log growth phase was extracted with the Ezup column bacteria genomic DNA purification kit and the Ezup column yeast genomic DNA purification kit before storage at −20°C for later use. The purity of genomic DNA was determined by the ratio of the absorbance at 260 nm to that at 280 nm (A_260_/A_280_) by an ultra-trace UV-visible spectrophotometer (Nanovue Plus, General Electric Co., MA, USA). The ratios of A_260_/A_280_ of genomic DNA from three strains were all greater than 1.8, which suggests that these DNA solutions were free from proteins (44). The interaction between GMM and genomic DNA was investigated by Liu, et al.’s method with some modification (45). The DNA solutions were diluted in sterile Tris-HCl to a final concentration of 3.6 mM, which was calculated by A_260_ in 1 cm quartz cell divided by a molar absorption coefficient ε_260_=6600M^-1^cm^-1^ (46). DNA diluents (250 μL) from *E. coli, S. aureus, and C. albicans* were added to 33 sterile 1.5 mL centrifuge tubes, and then various concentrations of GMM were then transferred to these tubes to reach final concentrations of 2, 4, 8, 16, 32, 64, 125, 250, 500, and 1000 μg/mL. The same volume of ethanol was added to the control groups. All mixtures were stirred upside down and allowed to equilibrate for 5 min. The UV spectral scanning was performed on a UV-visible spectrophotometer (Lambda 35, PerkinElmer Co., MA, USA) equipped with a xenon lamp. To eliminate the adverse effect from the background, the baseline was firstly corrected for Tris-HCl buffer signal before determination. The UV absorption spectra were recorded at a wavelength region ranging from 220–380 nm and measured in triplicate.

### Cell cycle analysis

**The** assay was conducted by flow cytometry combined with PI staining for DNA (47). Aliquot cell suspensions of 950 μL (cell density≈ 10^5^ CFU/mL) were prepared, and then to the suspension, 50 μL different concentrations of GMM were added to achieve final concentrations of 1/4 MIC, 1/2 MIC, 1 MIC, 2 MIC, and 4 MIC, respectively. Rather than GMM, 50 μL of ethanol was added to each of the control groups for the three microorganisms. All concentrations were performed three times in parallel. The mixed solutions were incubated for 1 h at 37°C (2 h at 28°C for *C. albicans*) with shaking at 120 rpm. Then, the cell pellets were collected by centrifugation and washed twice with sterile PBS. To the cell pellets, 70% (v/v) ice ethanol (after pre-cooling at −20°C overnight) was added to achieve fixation overnight at 4°C. Subsequently, the fixed cells underwent centrifugation and a PBS wash to eliminate the negative influence of the fixing fluid. Finally, the cell pellets were stained with 1 mL of 50 μg/mL PI solution (containing 1 mg/mL Rnase) and were let to stand for 20min at 4°C in darkness followed by flow cytometer detection.

**The** cell cycles were analyzed by following the process in the Beckman flow cytometer operation manual using ECD channel (610/20). The excitation and emission fluorescence wavelength of PI-bound DNA was located at 488 nm and 610 nm respectively. The sample flow rates were adjusted to 100–500 cells/s, and a minimum of 30,000 cells were collected for subsequent data processing.

### Real-time measurement of intracellular DNA, RNA, and protein content

**Three** dyes, Ho, PY, and FITC, were used to stain the DNA, RNA, and proteins, respectively, in cells. The amount of DNA, RNA, and protein was indirectly characterized by the light intensity of blue, red, and green fluorescence, respectively which had little overlap in the emission spectrum area (48). The 950 μL aliquot cell cultures (cell density≈ 10^5^ CFU/L) were treated with 50 μL GMM solution to reach final concentration of MIC. Similarly, 50 μL of ethanol were added to the control groups. All experiments were performed in triplicate. The cells were incubated 0, 10, 20, 30, 40, 50, and 60 min at 37°C (28°C for *C. albicans*) with shaking at 120 rpm. After different lengths of time, microbial cells were harvested by centrifugation and quickly transferred to 70% (v/v) ice ethanol (pre-cooled at −20°C overnight) followed by refrigeration at 4°C overnight to achieve cell fixation. On the second day, fixed cells underwent centrifugation and PBS wash to remove the ethanol. Subsequently, 1 mL of mixed dye work solution containing 0.5 μg/mL HO, 2.0 μg/mL PY and 0.1 μg/mL FITC was added to cells and allowed to stand for 20 min at 4°C in dark conditions. Finally, the concentration of stained cells was adjusted to 10^5^ CFU/mL with sterile PBS which ensured sufficient dyes for binding DNA, RNA, and protein prior to flow cytometry analysis.

**A** Beckman CytoFLEX flow cytometer equipped with three-laser excitation flow system was used to detect different fluorescence intensities. The excitation laser wavelengths of HO-DNA, PY-RNA, and FITC-protein were located at 355, 530, and 457 nm, respectively. Correspondingly, the emitted fluorescence intensities of HO-bound DNA (blue), PY-bound RNA (red), and FITC-bound protein (green) were measured at the wavelengths of 450, 580, and 520 nm respectively.

### Statistical analysis

**All** data were expressed as the means ± standard deviations (SD) in three replicate determinations. A multiple t test in GraphPad Prism 6 was used to analyze errors. A statistical significant difference exists if p<0.05.

## Acknowledgments

**This** work was supported by the National Key Technology R&D Program of China (No. 2017YFC1601000). **We** thank Cheng Hao for his assistance in using flow cytometer and Tao Ruan for his help in operating the scanning electron microscope. We thank the South China Institute of Collaborative Innovation for their support with bacteria and yeast genomic DNA extraction. The authors declare no competing financial interest.

